# mRNA-1273 vaccination protects against SARS-CoV-2 elicited lung inflammation in non-human primates

**DOI:** 10.1101/2021.12.24.474132

**Authors:** Adam T. Waickman, Kaitlin Victor, Krista Newell, Tao Li, Heather Friberg, Kathy Foulds, Mario Roederer, Diane L. Bolton, Jeffrey R. Currier, Robert Seder

## Abstract

Vaccine-elicited SARS-CoV-2 antibody responses are an established correlate of protection against viral infection in humans and non-human primates. However, it is less clear that vaccine-induced immunity is able to limit infection-elicited inflammation in the lower respiratory tract. To assess this, we collected bronchoalveolar lavage fluid samples post-SARS-CoV-2 strain USA-WA1/2020 challenge from rhesus macaques vaccinated with mRNA-1273 in a dose-reduction study. Single-cell transcriptomic profiling revealed a broad cellular landscape 48 hours post-challenge with distinct inflammatory signatures that correlated with viral RNA burden in the lower respiratory tract. These inflammatory signatures included phagocyte-restricted expression of chemokines such as *CXCL10* (IP10) and *CCL3* (MIP-1A) and the broad expression of interferon-induced genes such as *MX1, ISG15*, and *IFIT1*. Induction of these inflammatory profiles was suppressed by prior mRNA-1273 vaccination in a dose-dependent manner, and negatively correlated with pre-challenge serum and lung antibody titers against SARS-CoV-2 spike. These observations were replicated and validated in a second independent macaque challenge study using the B.1.351/beta-variant of SARS-CoV-2. These data support a model wherein vaccine-elicited antibody responses restrict viral replication following SARS-CoV-2 exposure, including limiting viral dissemination to the lower respiratory tract and infection-mediated inflammation and pathogenesis.

**One Sentence Summary:** Single cell RNA sequencing analysis demonstrates that mRNA-1273 vaccination limits the development of lower respiratory tract inflammation in SARS-CoV-2 challenged rhesus macaques

## INTRODUCTION

Severe acute respiratory syndrome coronavirus 2 (SARS-CoV-2) - the causative agent of COVID-19 – has infected at least 250 million individuals and resulted in over 5 million deaths as of November, 2021 (*1*). SARS-CoV-2 infection results in a range of clinical outcomes, from asymptomatic clearance to severe lung pathology with concomitant acute respiratory distress. Almost all morbidity and mortality attributable to SARS-CoV-2 is seen in the minority of patients who develop severe pneumonia requiring mechanical ventilation (*2, 3*). This has led to speculation that SARS-CoV-2 infection may promote a unique pathophysiology in which dysregulated immune responses to infection in the lower respiratory tract augment the severity of COVID-19. Indeed, examinations of the cellular composition of bronchoalveolar lavage fluid (BALF) from acutely ill COVID-19 patients have revealed a cellular landscape containing both resident cells and infiltrating immune cells displaying a unique and dysregulated inflammatory profile (*4*). Taken together, these data are consistent with a model wherein SARS-CoV-2 infected cells engage in a positive feedback loop with infiltrating immune cells to potentiate persistent alveolar inflammation and pathology (*5*). An effective vaccine would then be expected to impede the initiation of this, or a similar, pathological feedback loop thereby limiting lower airway disease.

Single cell RNA sequencing (scRNAseq) is a highly sensitive tool for analyzing the spectrum of SARS-CoV-2 elicited inflammation and the impact of vaccine-mediated immunity. The use of this approach to examine human BALF and PBMC samples has already identified several populations of immune cells likely implicated in inflammation-driven immunopathology and vaccine-mediated protection, as well as those likely to contain SARS-CoV-2 genetic material (*5-10*). Studies of human PBMC using scRNAseq have demonstrated a dysregulated response in both innate and adaptive immune cells in severe disease (*11*), evidence of emergency myelopoiesis cell and neutrophil dysregulation in severe disease (*6*), and an up-regulation of the TNF/IL-1β-driven inflammatory response as compared to influenza in classical monocytes (*12*). The unifying theme of these studies is that in severe COVID-19, compared to mild disease or asymptomatic infection, there is a profound and dysregulated type I interferon response across many lymphoid and myeloid origin cells (*13*). This response is accompanied by hyper-inflammation, evidence of cellular proliferation, and defective antigen-presentation and interferon responsiveness in classical monocytes.

Animal models have recapitulated many key aspects of the inflammatory response observed in the human lung, such as viral shedding, cellular infiltration profiles and cellular inflammatory profiles at the transcriptional level (*14, 15*). These models provide the critical ability to control dose and exposure variables that present a fundamental barrier to the accurate interpretation of human studies. Additionally, rhesus macaques *(Macaca mulatta)* represent a clinically relevant model for assessing lung tissue pathology and temporal analysis of SARS-CoV-2 elicited inflammation, and are the preclinical gold standard for assessing SARS-CoV-2 vaccine efficacy (*16-22*). Treatment of rhesus macaques with a clinically-approved JAK1/JAK2 inhibitor resulted in reduced lung inflammation and pathology, corresponding with attenuated infiltration of inflammatory immune cells and NETosis (*15*). These outcomes were associated with suppression of neutrophil recruitment and production of cytokines and chemokines by inflammatory macrophages, despite comparable type I IFN responses. Similarly, responses to SARS-CoV-2 in ferrets revealed a shift in BALF macrophage gene expression signatures toward a pro-inflammatory phenotype during early infection (*12*), underscoring the critical need to understand the cellular complexities of SARS-CoV-2 elicited inflammation.

mRNA-based vaccine platforms – such as Moderna’s mRNA-1273 and Pfizer/BioNTech’s BNT162b2 – which encode a stabilized version of the SARS-CoV-2 spike glycoprotein (*23*) show >90% efficacy against symptomatic COVID-19 in initial Phase 3 analyses and in large-scale prospective studies performed after their global rollout (*24, 25*). However, the efficacy of these vaccines against severe lower airway disease wanes over time after the initial prime and boost (*26-28*). Pre-clinical and clinical studies have strongly suggested that vaccine-elicited serum levels of SARS-CoV-2 neutralizing antibody titers are a mechanistic immune correlate of vaccine efficacy (*29, 30*). Despite the abundance of clinical and pre-clinical efficacy data for these mRNA-based vaccine platforms, there is little prospective information currently available on how these vaccines impact SARS-CoV-2-elicited inflammation in the lower respiratory tract with any degree of spatial or temporal resolution.

In this study we sought to understand the impact of mRNA-1273 vaccination on the cellular inflammatory response to SARS-CoV-2 infection in the lower respiratory tract of nonhuman primates (NHPs,) and whether vaccination is capable of breaking the inflammatory feedback loop that characterizes severe COVID-19. We used scRNAseq to analyze BALF cells from rhesus macaques challenged with SARS-CoV-2 strain USA-WA1/2020 after vaccination with two doses of 30μg or 1μg of mRNA-1273 or PBS. mRNA-1273 vaccination limited SARS-CoV-2 elicited inflammation in the lower respiratory tract as defined by the expression of pro-inflammatory chemokines and cytokines in multiple cell types, as well as the broad reduction in expression of interferon gene products such as *MX1, ISG15*, and *IFIT1*. Additionally, SARS-CoV-2 elicited inflammation was directly associated with post-challenge viral titers and inversely associated with pre-challenge antibody levels in unvaccinated and mRNA-1273 vaccinated animals. The ability of mRNA-1273 to limit SARS-CoV-2 elicited inflammation in the lower reparatory tract was independently verified using the antigenically disparate B.1.351/beta variant. Collectively, these results demonstrate that vaccination with mRNA-1273 not only limits SARS-CoV-2 viral replication, but restricts inflammation in NHPs. Additionally, these data support a model wherein neutralizing antibody at the site of virus inoculation reduces the viral burden, constraining upper respiratory tract viral replication and secondary viral dissemination to the lower respiratory tract and infection-associated inflammation.

## RESULTS

### Frequency of BALF resident cells following SARS-CoV-2 challenge

It has previously been demonstrated that vaccination of macaques with mRNA-1273 results in robust serum antibody responses and high-level protection from subsequent SARS-CoV-2 challenge in a dose-dependent fashion (*19, 29*). To extend these observations and to assess the impact of vaccination on SARS-CoV-2 elicited inflammation in the lower respiratory tract, we performed scRNAseq analysis of fresh BALF obtained on days 2 and 7 post SARS-CoV-2 challenge in animals which previously received either 30 μg (n=4) or 1 μg (n=6) of mRNA-1273 in a prime-boost series administered four weeks apart. Control animals received PBS (n=6). All animals were challenged intranasally/intratracheally (IN/IT) with 8×10^5^ PFU of SARS-CoV-2 (strain USA-WA1/2020) four weeks after the last vaccine dose. In addition, BALF cells were analyzed from naïve uninfected animals to serve as controls.

scRNAseq was used to classify and quantify the cell composition and dynamics within the BALF post-challenge. A total of 65,226 viable and high quality BALF cells from all animals were recovered after filtering and quality control steps (**Fig. 1A, 1B**). Of note, epithelial cells (**Fig. 1C**), lymphocytes (**Fig. 1D**), dendritic cells (**Fig. 1E**) and macrophages (**Fig. 1F**) were identified in all time points from all animals. Alveolar macrophages were further separated into either MARCO^-^ or MARCO^+^ populations, corresponding to interstitial and tissue-resident alveolar macrophages, respectively (*14, 31*). Following SARS-CoV-2 challenge, CD4^+^ and CD8^+^ T cells increased in frequency between days 2 and 7 post-challenge in unvaccinated animals. Several DC populations also trended higher among unvaccinated infected animals at one or more time points relative to uninfected controls. No significant changes were observed in the frequency of epithelial cell or macrophage populations.

**Fig 1.**
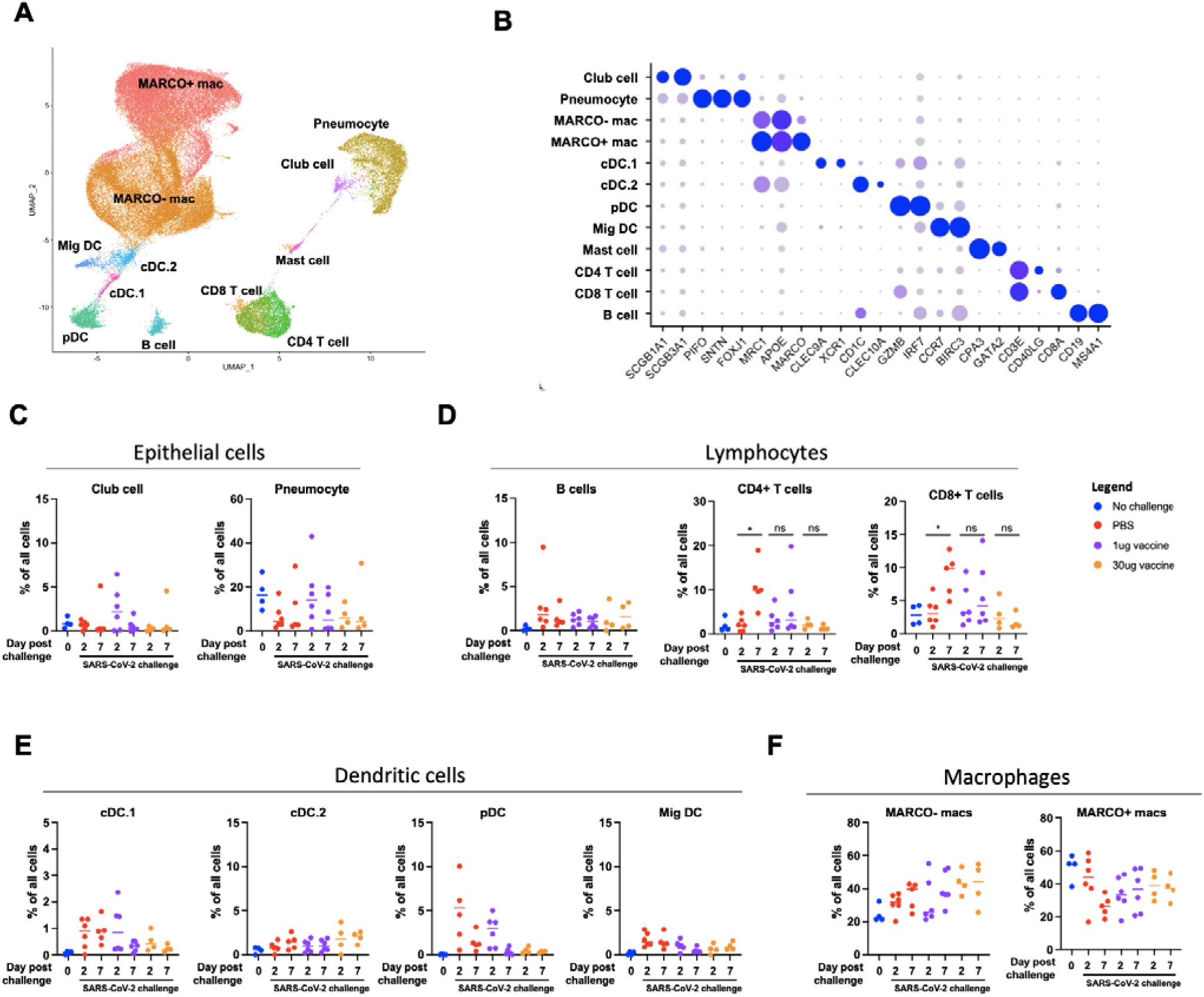
Identification and quantification of BALF cells by scRNAseq. **A)** UMAP projection of BALF cells captured by scRNAseq analysis. **B)** Expression of key linage specific genes in all annotated cell types. **C)** Frequency of epithelial cell populations. **D)** Frequency of lymphocyte cell populations. **E)** Frequency of dendritic cell populations. **F)** Frequency of macrophage populations

### Inflammatory signatures of SARS-CoV-2 infection

To assess the inflammatory response elicited by SARS-CoV-2 challenge in naïve and mRNA-1273 vaccinated animals, the expression of inflammatory markers, chemokines, and cytokines was assessed in each annotated BALF cell type on days 2 and 7 post challenge. An inflammatory response to infection – as indicated by the expression of genes such as *MX1, ISG15*, and *IFIT1* – was observed across all cell types in the unvaccinated infected animals on day 2 post SARS-CoV-2 challenge. (**Fig. 2A**). Expression of these markers decreased in a dose-dependent fashion in animals vaccinated with 1 μg or 30 μg of mRNA-1273. These transcriptional signatures of acute viral infection resolved in most cell types by day 7 post infection, with the exception of lingering *MX1/MX2* expression in some populations of macrophages and DCs (**Fig. 2B)**. Migratory DCs and MARCO^-^ macrophages responded to SARS-CoV-2 challenge in unvaccinated animals by expressing chemokines such as *CXCL10* (IP10) an *CCL3* (MIP-1A), both of which were previously identified in the context of acute SARS-CoV-2 infection in humans (*4*). In addition, elevated expression of cytotoxic factors *GZMA* and *PRF1* was observed in CD8^+^ T cells following SARS-CoV-2 challenge on day 2 and maintained 7 days post challenge. Notably, the expression of these pro-inflammatory chemokines and chemokines was dramatically suppressed in vaccinated animals in a dose-dependent manner across all time points.

**Fig 2.**
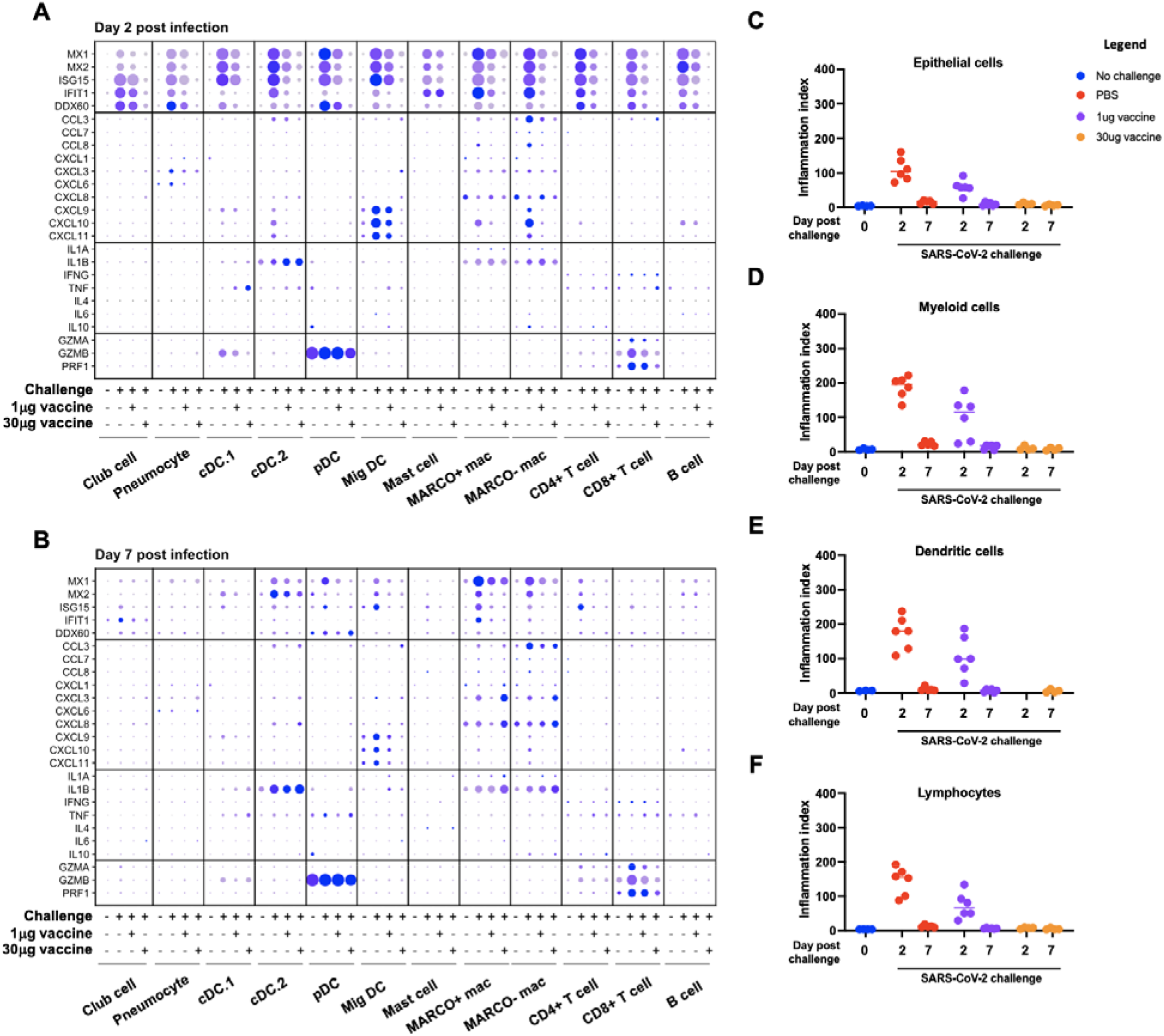
Transcriptional signatures of SARS-CoV-2 WA-1 elicited inflammation. **A)** Expression of inflammatory markers and cytokines/chemokines in all annotated cells day 2 post infection. **B)** Expression of inflammatory markers and cytokines/chemokines in all annotated cells day 7 post infection. **C)** Inflammatory index scores in epithelial cells. **D)** Inflammatory index scores in dendritic cells. **E)** Inflammatory index scores in myeloid cells. **F)** Inflammatory index scores in lymphocytes

To reduce the complexity of the data and provide more direct insight into the dynamics of SARS-CoV-2 elicited inflammation, we defined a transcriptional “inflammation index” which could be used to quantify the level of enrichment for inflammatory gene products in a given sample and cell type. This index was developed by selecting 8 genes (*MX1, MX2, IFIT1, IFIT2, IFIT3, IFI6, ISG15*, and *ISG20*) that were 1) previously known to be regulated at a transcriptional level by viral infection and/or interferon stimulation, 2) highly induced in our dataset following SARS-CoV-2 challenge, and 3) consistently observed in all cell types captured in our analysis. Using this reductionist approach, we observed a dose-dependent suppression of SARS-CoV-2 elicited inflammation in epithelial cells (pneumocytes, club cells), myeloid cells (MARCO^+^ macrophages, MARCO^-^ macrophages, mast cells), dendritic cells (cDC.1, cDC.2, pDC, Mig DC), and lymphocytes (B cells, CD8^+^ T cells, CD4^+^ T cells) with increasing mRNA vaccination dose (**Fig. 2C-F**). Furthermore, inflammation in animals that received the full dose of vaccine was nearly equivalent to that of the unchallenged control animals in all cell types assessed. Inflammation returned to baseline in all groups by day 7 post vaccination. These results establish a single metric for quantifying the transient inflammatory transcriptional response elicited following SARS-CoV-2 infection across multiple cell populations using scRNAseq, and by extension provide a measurement of the site-specific host-response to the virus.

### Cell-associated viral RNA burden following SARS-CoV-2 infection

Having defined the impact of mRNA-1273 vaccination on SARS-CoV-2 associated inflammation in the lower respiratory tract of macaques, we next attempted to quantify the cell-associated SARS-CoV-2 viral RNA burden by aligning scRNAseq reads that failed to align to the macaque genome against the SARS-CoV-2 USA-WA1/2020 reference genome. BALF contained widespread SARS-CoV-2 RNA^+^ cells on day 2 in unvaccinated animals (**Fig. 3A, 3B**). SARS-CoV-2 RNA^+^ positive cells were seen in all annotated cell types with the exception of mast cells in unvaccinated animals, although the greatest number of viral RNA^+^ cells were found in the MARCO^-^ macrophage cluster. Similar to the inflammation index, the frequency of viral RNA^+^ cells was suppressed by vaccination in a dose-dependent fashion and mostly resolved by day 7 post infection. The frequency of viral RNA^+^ cells in the BALF on day 2 correlated well with contemporaneous viral subgenomic RNA (sgRNA) load in the BALF as quantified by PCR of the E and N gene (**Fig. 3C, fig. S1**). Notably, the correlation between the frequency of viral RNA^+^ cells in the BALF was weaker with the upper respiratory tract (nasopharyngeal swab) sgRNA loads (**Fig. 3D, fig. S1**). These results show that SARS-CoV-2 viral burden in lung cells is abrogated by mRNA vaccination and is consistent with reduced soluble viral RNA measures in BALF.

**Fig 3.**
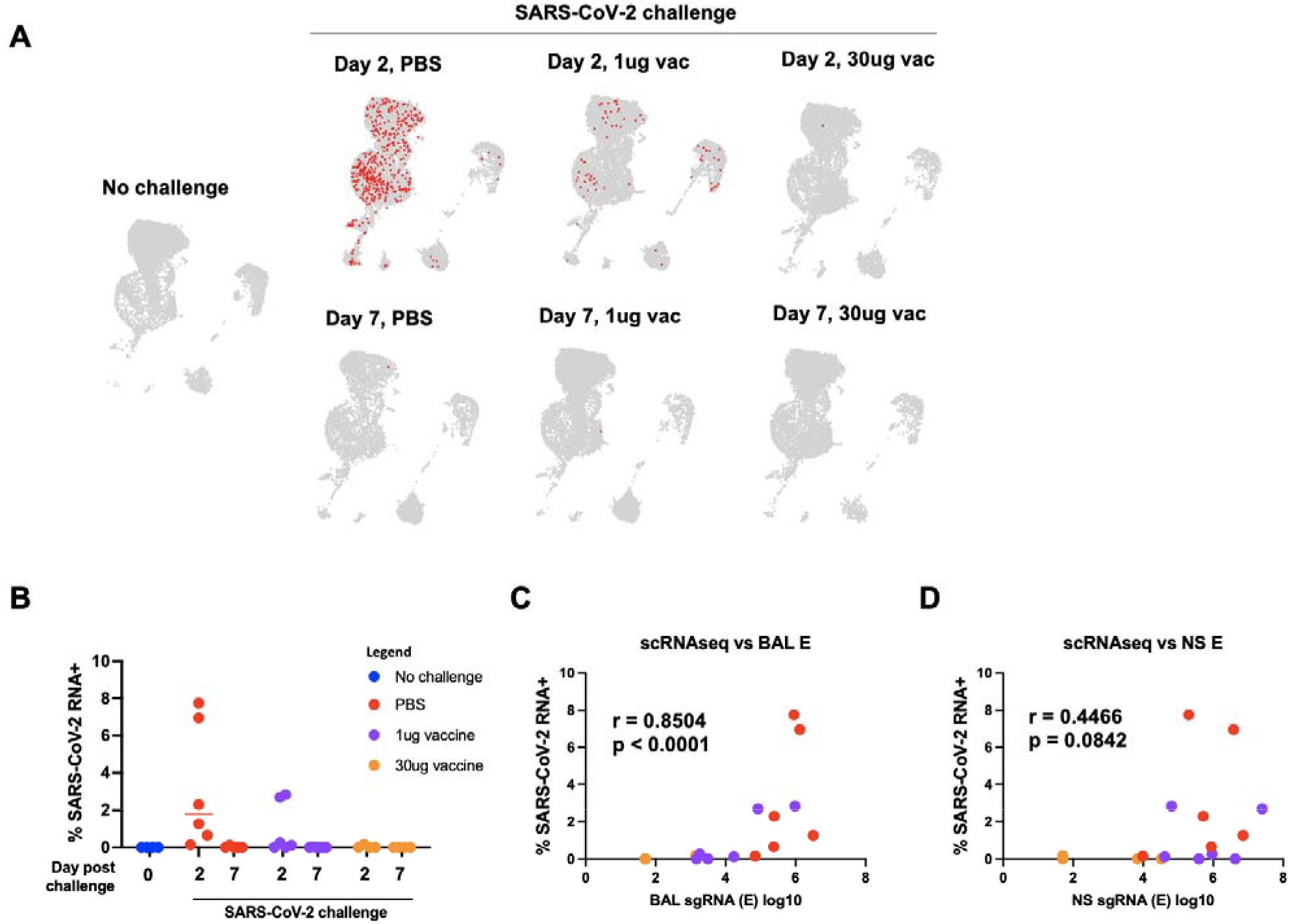
Identification and quantification of SARS-CoV-2 RNA positive cells. **A)** Location of SARS-CoV-2 RNA positive cells. **B)** Frequency of SARS-CoV-2 RNA positive cell. **C**) Relationship between BALF RNA load and frequency of SARS-CoV-2 RNA^+^ cells. **D)** Relationship between NS RNA load and frequency of SARS-CoV-2 RNA^+^ cells. Spearman correlation.

### Relationship between SARS-CoV-2 RNA load and cell type-specific inflammation

To examine the relationship between the observed dose-dependent reduction in SARS-CoV-2 viral burden and inflammation in the BALF of mRNA-1273 vaccinated animals after SARS-CoV-2 challenge, we compared viral RNA measures to cellular inflammatory responses. Strikingly, cell-free SARS-CoV-2 RNA load positively correlated with the previously defined inflammation index score of both BALF dendritic and myeloid cell compartment on day 2 post infection across all study groups (**Fig. 4A, Fig. 4D, fig. S2**). However, nasal swab viral RNA load poorly correlated with dendritic cell inflammation, and correlated only weakly with myeloid inflammation (**Fig. 4B, Fig. 4E, fig. S2**). Cell-associated viral RNA loads in the BALF also correlated with the inflammation score for both dendritic and myeloid compartments (**Fig. 4C, Fig. 4F**). These results demonstrate that viral burden in the BALF, but not nasal environment, correlates with the amount of lower respiratory tract inflammation following SARS-CoV-2 challenge.

**Fig 4.**
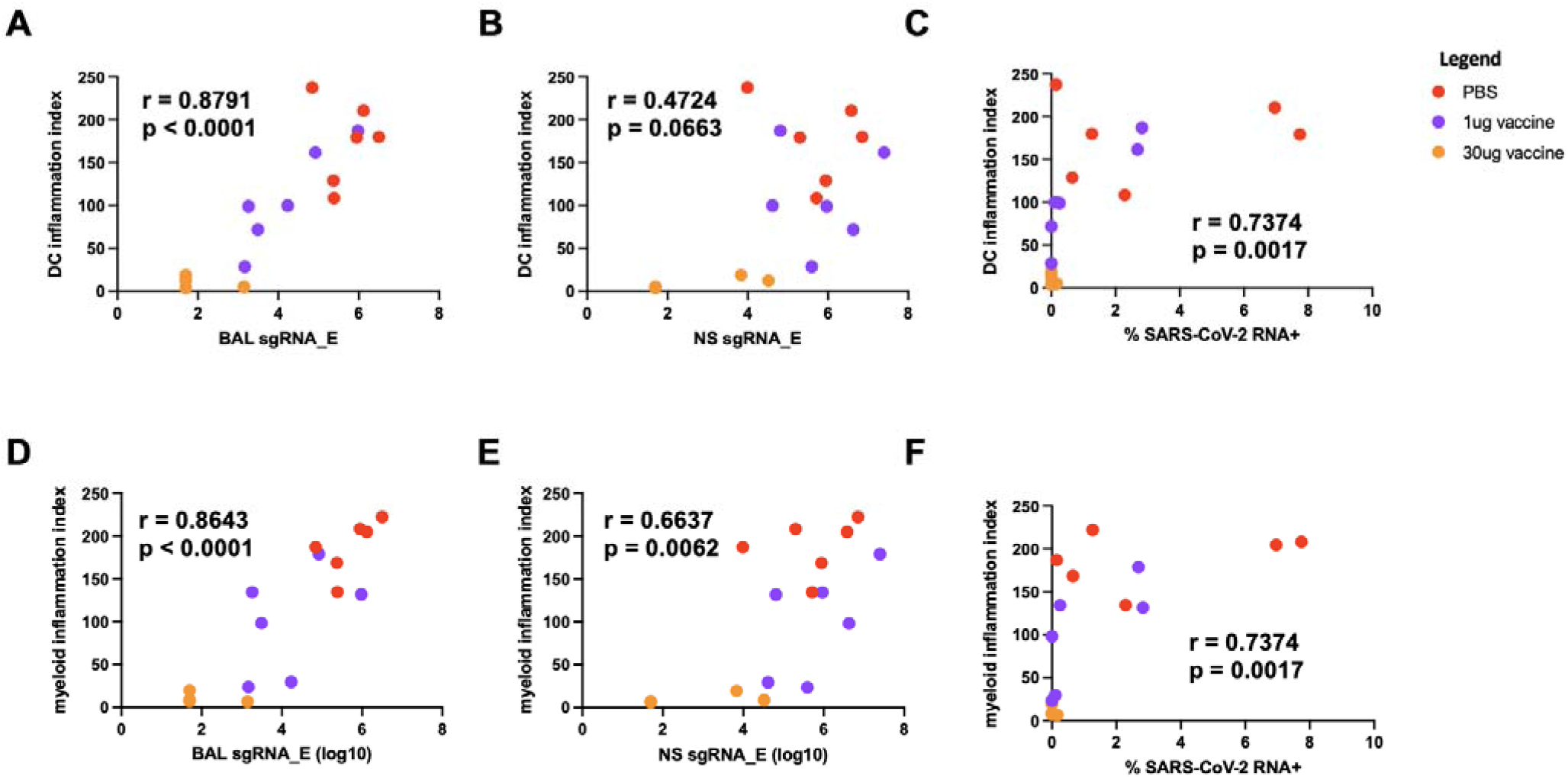
Viral load vs inflammation day 2 post infection. **A)** DC Inflammation index score vs BALF sgRNA (E gene). **B)** DC Inflammation index score vs Nasal swab sgRNA (E gene). **C)** DC Inflammation index score vs SARS-CoV-2 RNA^+^ cell fraction. **D)** Macrophage Inflammation index score vs BALF sgRNA (E gene). **E)** Macrophage Inflammation index score vs Nasal swab sgRNA (E gene). **F)** Macrophage Inflammation index score vs SARS-CoV-2 RNA^+^ cell fraction. Spearman correlation.

### Pre-challenge immune profiles predict lung inflammation following SARS-CoV-2 challenge

SARS-CoV-2-specific antibody titers have been implicated in mRNA vaccination-mediated protection from SARs-CoV-2 infection in both humans and NHPs (*29, 30, 32*), but the relationship between specific antibody titers and lower respiratory tract inflammation is not clear. To this end, we incorporated previously published data (*29*) on serum levels of full-length spike protein and receptor-binding domain (RBD)-specific titers IgG present immediately before SARS-CoV-2 challenge in these animals into our analysis. Pre-challenge (8 week post initial vaccine dose) serum titers of both spike- and RBD-specific IgG were negatively correlated with dendritic cell inflammation scores in the lower respiratory tract 2 days post SARS-CoV-2 challenge across all study groups (**Fig. 5A to B**). This relationship was also observed with pre-challenge spike-specific IgG titers in the BALF (**Fig. 5C)**. Furthermore, serum neutralizing antibody responses assessed by both pseudovirus and live-virus neutralization assays were also associated with reduced DC inflammatory responses (**Fig. 5D to E**). These data suggest that SARS-CoV-2 specific antibody titers in mRNA-1273 vaccinated macaques function as a powerful predictor of SARS-CoV-2 elicited inflammation in the lower respiratory tract.

**Fig 5.**
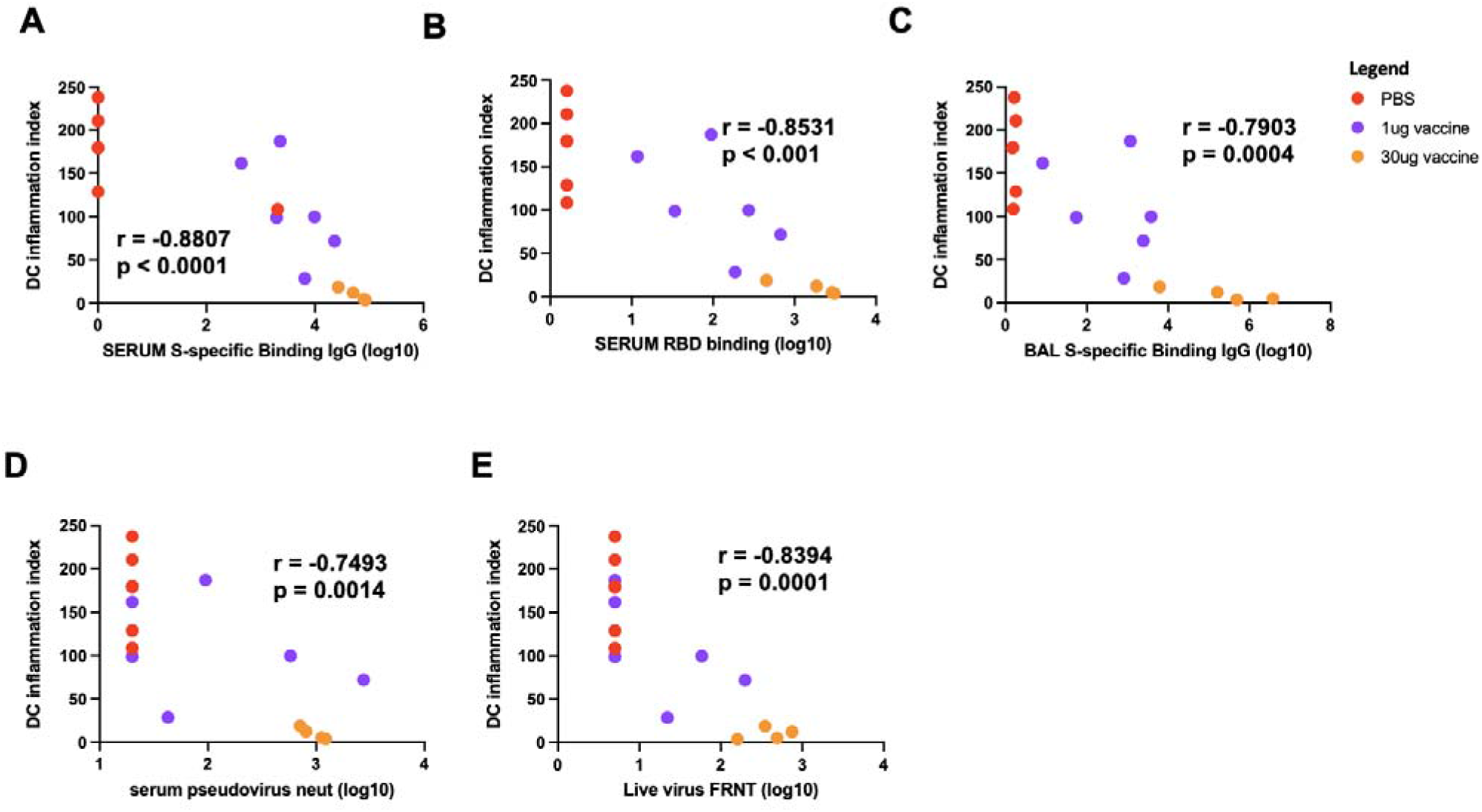
Relationship between antibody titers and SARS-CoV-2 elicited inflammation. **A)** Relationship between pre-challenge serum S-specific IgG titers (wk 8 post vaccination) and DC inflammation on day 2 post challenge. **B)** Relationship between pre-challenge serum RBD-specific IgG titers (wk 8 post vaccination) and DC inflammation on day 2 post challenge. **C)** Relationship between pre-challenge BALF S-specific IgG titers (wk 6 post vaccination) and DC inflammation on day 2 post challenge. **D)** Relationship between pre-challenge serum pseudovirus neut titers titers (wk 8 post vaccination) and DC inflammation on day 2 post challenge. **E)** Relationship between pre-challenge serum live virus FRNT (wk 8 post vaccination) and DC inflammation on day 2 post challenge. Spearman correlation

### Impact of mRNA-1273 vaccination on inflammation elicited by SARS-CoV-2 B.1.351/beta variant

Having established the lower respiratory tract profile associated with SARS-CoV-2 USA-WA1/2020 infection – and how prior mRNA-1273 immunization blunts infection-attendant inflammation in this macaque model – we sought to expand and validate our observations in an independent experiment using a SARS-CoV-2 variant challenge. We again utilized scRNAseq to analyze BALF resident cells isolated on day 2 post challenge with the SARS-CoV-2 B.1.351/beta variant in naïve animals, or animals vaccinated twice with 30μg of mRNA-1273. The same populations of BALF resident cells identified following WA-1 challenge were observed following B.1.351/beta challenge (**Fig. 6A, fig. S3**). However, unlike USA-WA1/2020 challenge, infection with B.1.351/beta resulted in a significant perturbation in the abundance of multiple cell types including pDCs and migratory DCs (**fig. S4**). These changes in cellularity were not observed in mRNA-1273 vaccinated animals, and the production of chemokines, cytokines, and cytolytic factors were again suppressed in vaccinated animals relative to their unvaccinated counterparts (**Fig. 6B**). Vaccination with mRNA-1273 also suppressed SARS-CoV-2 associated inflammation observed in epithelial cells (**Fig. 6C**), dendritic cells (**Fig. 6D**), myeloid cells (**Fig. 6E**), and lymphocytes (**Fig. 6F**). The frequency of SARS-CoV-2 RNA positive cells in BALF was also reduced by mRNA-1273 vaccination, with the greatest number of viral RNA^+^ cells again found in the MARCO^-^ macrophage cluster (**fig. S5**). In their totality, these results indicate that vaccination with mRNA-1273 is capable of limiting lower airway inflammation in macaques following challenge with multiple antigenically and evolutionally divergent strains of SARS-CoV-2.

**Fig 6.**
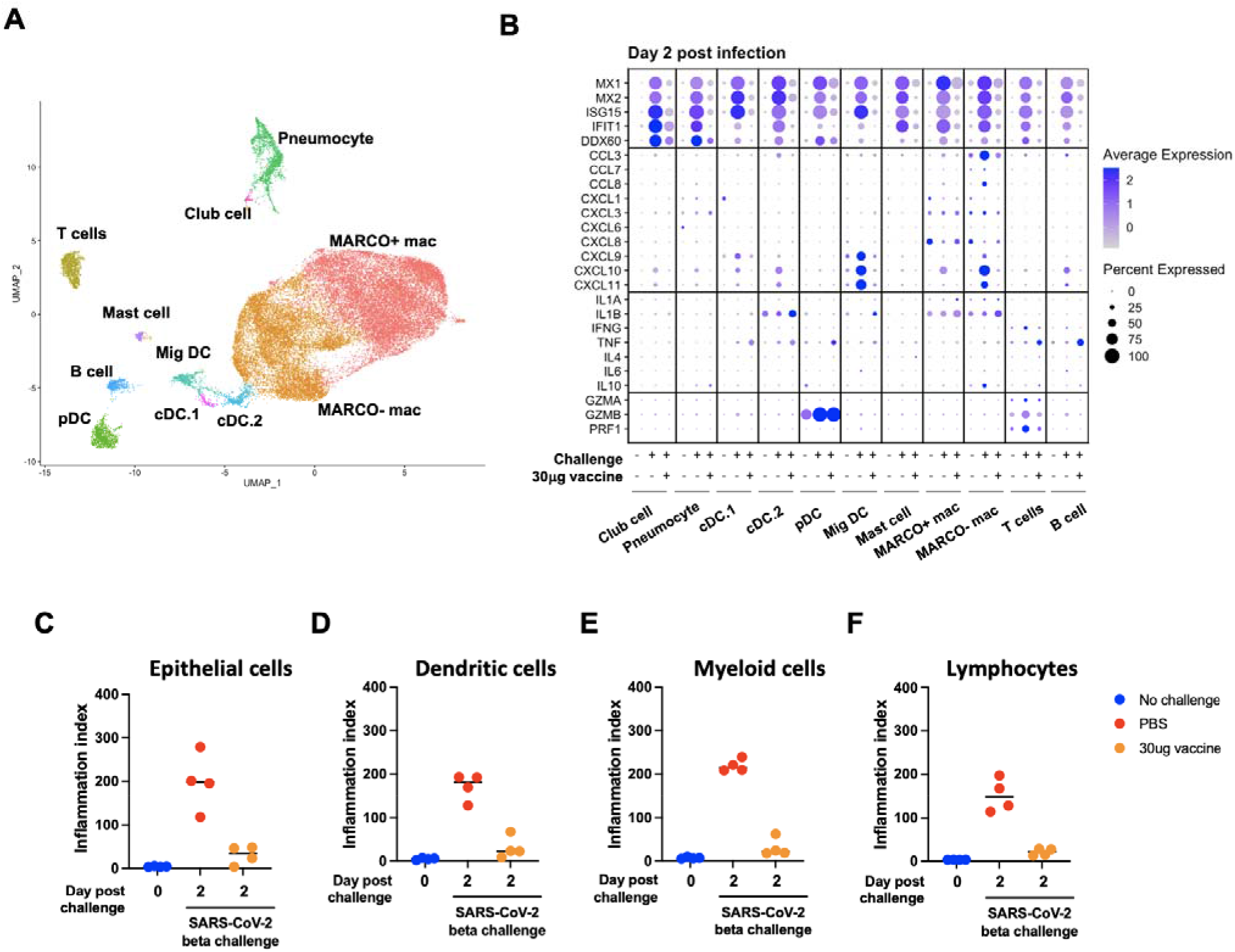
Transcriptional signatures of SARS-CoV-2 beta VBM elicited inflammation. **A)** UMAP projection of BALF cells from beta VBM challenge. **B)** Expression of inflammatory markers and cytokines/chemokines in all annotated cells day 2 post beta infection. **C)** Inflammatory index scores in epithelial cells. **D)** Inflammatory index scores in dendritic cells. **E)** Inflammatory index scores in myeloid cells. **F)** Inflammatory index scores in lymphocytes

## DISCUSSION

In this study we sought to provide functional and mechanistic insight into the properties of mRNA-1273 elicited protection from SARS-CoV-2 challenge in a widely used nonhuman primate model of mild to moderate COVID-19 disease. While immune correlates of protection from symptomatic SARS-CoV-2 infection are currently being assessed and defined in both clinical and pre-clinical studies, there is a more limited information on the impact of vaccine-elicited immunity on SARS-CoV-2 induced inflammation in the lungs at the single cell level. Here, we utilized scRNAseq technology to analyze BALF cells from mRNA-1273 vaccinated animals that were subsequently challenged with SARS-CoV-2 to define the transcriptional signatures of infection and to ascertain how this inflammatory response is modulated by vaccine-elicited adaptive immune responses in a dose-dependent fashion. SARS-CoV-2 infection induced a robust inflammatory response in all unvaccinated animals that was suppressed in a dose-dependent fashion by mRNA-1273 vaccination. Notably, migratory DCs and MARCO^-^ macrophages appeared to be the most responsive cell types in the lower respiratory tract to SARS-CoV-2 infection, as indicated by chemokine and cytokine production. Cell-associated SARS-CoV-2 viral RNA was readily detected in the BALF of unvaccinated animals, restricted by mRNA-1273 vaccination, and correlated with dendritic and myeloid cell inflammation.

It was previously established that S-specific antibody responses elicited by mRNA-1273 vaccination correlates with upper and lower airway control of SARS-CoV-2 replication in macaques after challenge (*29*). Here we further demonstrate, in these same animals, that the pre-challenge antibody profile, including titers of binding and neutralizing antibody, predicted and inversely correlated with the inflammatory profile within the lung across multiple cell types. The high degree of correlation between pre-challenge antibody titers, post-challenge viral loads, and post-challenge inflammation suggests a model of mRNA-1273-mediated protection from SARS-CoV-2 challenge in nonhuman primates. Namely, neutralizing antibody levels determine the burden of viral replication, and the amount of virus persisting in the upper respiratory tract drives secondary viral dissemination to the lower respiratory tract and infection-attendant inflammation. The absence of inflammation and viral RNA in the lower respiratory tract of SARS-CoV-2 challenged animals just 2 days post infection supports the high level of efficacy of mRNA vaccination against lower respiratory infection and pathology.

Examination of human BALF from mild and severe COVID-19 patients has been used to distinguish an inflammatory signature associated with severity. This signature includes expression of genes for chemokine production, proinflammatory cytokines, and activated phenotypic markers within resident and infiltrating cells (*5-10*). Several studies have described the potential role of neutrophils and NETosis in local lung pathology, as well as shifts in monocyte and macrophage populations toward an inflammatory phenotype (*5-7, 9, 10*). Reinforcing the critical role of type I interferon in the antiviral response, many studies have also identified strong type I IFN signatures in single immune cells from COVID-19 patients, although the relationship of this cytokine profile and disease severity is still uncertain (*12, 33*). Accordingly, our observation of stronger correlation between BALF inflammatory immune cell gene signatures and BALF viral burden than that of nasopharyngeal swabs suggests that inflammation-driven lung pathology is directly influenced by local viral replication. However, given the migratory nature of these cell populations and the relatively low abundance of viral RNA in the lower respiratory tract, the possibility that these cells were stimulated by viral ligands at other anatomical sites cannot be discounted. Furthermore, the cells that were found to be positive for SARS-CoV-2 RNA represented a range of cell types. Although the range of cell types expressing ACE2 and therefore permissive to viral entry is wide (*34*), these populations may not represent bona fide productively-infected cells. Rather, cells may acquire viral RNA through phagocytic mechanisms, for example. Despite these potential caveats, our findings were validated by challenge of animals with the SARS-CoV-2 B.1.351/beta variant, wherein mRNA-1273 vaccination also prevented infection-induced lung inflammation.

There are some limitations of this study to consider. First, the relatively transient and self-limiting nature of SARS-CoV-2 infection and infection-elicited inflammation in macaques makes it difficult to place these observations into context of human disease. Many of the pathways, cell types, transcriptional signatures, and correlates of protection identified in our analysis have also been defined in humans with acute COVID-19, but the magnitude and timing of the events may not be homologous. Second, the route of virus administration has been shown to influence infection and inflammation in other models of SARS-CoV-2 challenge (*35*), so that the IN/IT route of infection used in this study may result in subtly different features of infection and inflammation than aerosol-mediated infection.

In conclusion, this study defines the lower respiratory tract cellular and transcriptional signature associated with SARS-CoV-2 infection in macaques using two distinct viral variants, and identifies conserved signatures of vaccine-elicited protection from infection-attendant inflammation. These data emphasize the contribution of inflammatory/migratory DCs and macrophages to lower respiratory tract inflammation following SARS-CoV-2 infection, and define a critical relationship between antibody titers, post-challenge viral burden, and broad infection-elicited inflammation.

## MATERIALS AND METHODS

### Vaccine formulation

mRNA encoding a sequence-optimized and prefusion-stabilized SARS-CoV-2 S-2P protein (*36, 37*) was synthesized in vitro and formulated as previously reported (*19, 38, 39*).

### Rhesus macaque vaccination model

3- to 8-year-old rhesus macaques of Indian origin were sorted by sex, age and weight and then stratified into groups as previously described (*29, 30*). Animals were immunized intramuscularly at week 0 and at week 4 with 1 μg or 30 μg of mRNA-1273 in 1 mL of PBS into the right hindleg. Placebo-control animals were administered an equal volume of PBS. Animal experiments were performed in compliance with all pertinent National Institutes of Health regulations and approval from the Animal Care and Use Committees of the Vaccine Research Center and Bioqual Inc. (Rockville, MD). Research was conducted under an approved animal use protocol in an AAALAC accredited facility in compliance with the Animal Welfare Act and other federal statutes and regulations relating to animals and experiments involving animals and adheres to principles stated in the Guide for the Care and Use of Laboratory Animals, NRC Publication, 2011 edition. Studies were conducted at Bioqual Inc. Post-vaccination antibody titers generated as previously described *(29, 38, 40-43)*, and previously reported by Corbett *et al*. (*29*).

### USA-WA1/2020 challenge

At week 8 post initial vaccination (4 weeks after boost), all animals were challenged with a total dose of 8 × 10^5^ PFUs of SARS-CoV-2 as previously described (*29*). The stock of 1.99 × 10^6^ TCID_50_ or 3×10^6^ PFU/mL SARS-CoV-2 USA-WA1/2020 strain (BEI: NR-70038893) was diluted and administered in 3-mL doses by the intratracheal route and in 1-mL doses by the intranasal route (0.5 mL per nostril). Post-challenge SARS-CoV-2 sgRNA burden in nasal swabs and BAL were determined as previously described (*19, 29*), and previously reported by Corbett *et al*. (*29*).

### B.1.351 challenge

At week 8 post initial vaccination (4 weeks after boost) NHPs were challenged with a total dose of 5×10^5^ PFU of SARS-CoV-2 B.1.351 strain as previously described (*30*). The viral inoculum was administered as 3.75×10^5^ PFU in 3 mL intratracheally (IT) and 1.25×10^5^ PFU in 1 mL intranasally (IN) in a volume of 0.5 mL into each nostril.

### scRNAseq library generation

Freshly isolated BALF suspensions were prepared for single-cell RNA sequencing using the Chromium Single-Cell 5′ Reagent v2 kit or NextGEM v1.0 kit and the Chromium Single-Cell Controller (10x Genomics, CA) (*44*). 2000–8000 cells per reaction suspended at a density of 50–500 cells/μL in PBS plus 0.5% FBS were loaded for gel bead-in-emulsion (GEM) generation and barcoding. Reverse transcription, RT-cleanup, and cDNA amplification were performed to isolate and amplify cDNA for downstream library construction according to the manufacturer’s protocol. Libraries were constructed using the Chromium Single-Cell 5′ reagent kit and i7 Multiplex Kit (10x Genomics, CA) according to the manufacturer’s protocol.

### Sequencing

scRNAseq 5′ gene expression libraries were sequenced on an Illumina NovaSeq 6000 instrument using the S1, S2, or S4 reagent kits (300 cycles). Libraries were balanced to allow for ∼150,000 reads/cell for 5′ gene expression libraries. Sequencing parameters were set for 150 cycles for Read1, 8 cycles for Index1, and 150 cycles for Read2. Prior to sequencing, library quality and concentration were assessed using an Agilent 4200 TapeStation with High Sensitivity D5000 ScreenTape Assay and Qubit Fluorometer (Thermo Fisher Scientific) with dsDNA BR assay kit according to the manufacturer’s recommendations.

### scRNAseq gene expression analysis/visualization

5′ gene expression alignment from all BALF samples was performed using the 10x Genomics Cell Ranger pipeline (*44*). Sample demultiplexing, alignment, barcode/UMI filtering, and duplicate compression was performed using the Cell Ranger software package (10x Genomics, CA, v2.1.0) and bcl2fastq2 (Illumina, CA, v2.20) according to the manufacturer’s recommendations, using the default settings and mkfastq/count commands, respectively. All reads were trimmed to 26bp x 98bp for gene expression analysis. Transcript alignment was performed against a *Macaca mulataa* reference library generated using the Cell Ranger mkref command, the Ensembl Mmul_10 top-level genome FASTA, and the corresponding Ensembl v100 gene GTF.

Multi-sample integration, data normalization, dimensional reduction, visualization, and differential gene expression were performed using the R package Seurat (v4.0.0) (*45, 46*). All datasets were filtered to only contain cells with between 200–5,000 unique features and <12.5% mitochondrial RNA gene content (defined as expression of the following mitochondrial gene products: *ND1, ND2, COX1, COX2, ATP8, ATP6, COX3, ND3, ND4L, ND4, ND5*, and *CYTB*). To eliminate erythrocyte contamination, datasets were additionally filtered to contain cells with less than a 10% erythrocytic gene signature (defined as HBA and HBB). Data were scaled, normalized, and transformed prior to multi-sample integration using the negative binomial regression model of the Seurat SCTransform() function, additionally regressing-out the contribution of imputed cell cycle to the normalized dataset (*47*).

SelectIntegrationFeatures() and PrepSCTIntegration() functions were used to identify conserved features for dataset integration, and final dataset anchoring/integration were performed using FindIntegrationAnchors() and IntegrateData() functions, with the day 2, 30μg vaccine samples used as reference datasets. PCA was performed using variable genes defined by SCTransform(). For the USA-WA1/2020 dataset, the first 40 resultant PCs were initially used to perform a UMAP dimensional reduction of the dataset (RunUMAP()) and to construct a shared nearest neighbor graph (SNN; FindNeighbors()). This SNN was used to cluster the dataset (FindClusters()) with default parameters and resolution set to 0.7. From this initial clustering a population of low-viability cells was identified and removed from the anaylsis, after which the dataset PCA was re-run and the first 35 resultant PCs were used to perform a UMAP dimensional reduction of the dataset (RunUMAP()) and to construct a shared nearest neighbor graph (SNN; FindNeighbors()). This SNN was used to cluster the dataset (FindClusters()) with default parameters and resolution set to 1.5. The resultant clusters were assigned to following cell types based on the expression of the indicated gene products: club cell *(SCGB1A1, SCGB3A1)*, pneumocyte (*PIFO, SNTN, FOXJ1*), MARCO^-^ mac (*MRC1, APOE*), MARCO^+^ mac (*MRC1, APOE, MARCO*), cDC.1 (*CLEC9A, XCR1*), cDC.2 (*CD1C, CLEC10A*), pDC (*GZMB, IRF7*), MigDC (*CCR7, BIRC3*), Mast cell (*CPA3, GATA2*), CD4^+^ T cell (*CD3E, CD40LG*), CD8^+^ T cell (*CD3E, CD8A*), and B cells (*CD19, MS4A1*).

For the B.1.351/beta dataset, the first 31 resultant PCs were initially used to perform a UMAP dimensional reduction of the dataset (RunUMAP()) and to construct a shared nearest neighbor graph (SNN; FindNeighbors()). This SNN was used to cluster the dataset (FindClusters()) with default parameters and resolution set to 1.7. From this initial clustering a population of low-viability cells was identified and removed from the anaylsis, after which the dataset PCA was re-run and the first 31 resultant PCs were used to perform a UMAP dimensional reduction of the dataset (RunUMAP()) and to construct a shared nearest neighbor graph (SNN; FindNeighbors()). This SNN was used to cluster the dataset (FindClusters()) with default parameters and resolution set to 1.7. The resultant clusters were assigned to following cell types based on the expression of the indicated gene products: club cell (*SCGB1A1, SCGB3A1*), pneumocyte (*PIFO, SNTN, FOXJ1*), MARCO^-^ mac (*MRC1, APOE*), MARCO^+^ mac (*MRC1, APOE, MARCO*), cDC.1 (*CLEC9A, XCR1*), cDC.2 (*CD1C, CLEC10A*), pDC (*GZMB, IRF7*), MigDC (*CCR7, BIRC3*), Mast cell (*CPA3, GATA2*), CD4^+^ T cell (*CD3E, CD40LG*), CD8^+^ T cell (*CD3E, CD8A*), and B cells (*CD19, MS4A1*).

Following dataset integration and dimensional reduction/clustering, gene expression data was log transformed and scaled by a factor of 10,000 using the NormalizeData() function. This normalized gene expression data was used to determine cellular cluster identity by utilizing the Seurat application of a Wilcoxon rank-sum test (FindAllMarkers()), and comparing the resulting differential expression data to known cell-linage specific gene sets. Differential gene expression analysis between study time points was performed using normalized gene expression data and the Wilcoxon rank-sum test with implementation in the FindMarkers() function, with a log_2_ fold change threshold of 0.5 and min.pct of 0.25. Bonferroni correction was used to control for False Discovery Rate (FDR), with a corrected *p* value of < 0.05 considered significant.

### Identification of SARS-CoV-2 RNA^+^ cells

Quantification and alignment of cell-associated SARS-CoV-2 RNA from was performed using the 10x Genomics Cell Ranger pipeline(*44*). Sample demultiplexing, alignment, barcode/UMI filtering, and duplicate compression was performed using the Cell Ranger software package (10× Genomics, CA, v2.1.0) and bcl2fastq2 (Illumina, CA, v2.20) according to the manufacturer’s recommendations, using the default settings and mkfastq/count commands, respectively. The resulting untrimmed (150bp x 150bp) FASTQs were filtered to only contain reads that did not align to the *Macaca mulataa* genome using seqfilter and the read annotation from the Cellranger alignment performed on the trimmed FASTQs performed against the Ensembl Mmul_10 genome described above. Transcript alignment was performed against a SARS-CoV-2 reference generated using the Cell Ranger mkref command and the SARS-CoV-2 reference genome (strain USA-WA1/2020) FASTA and the corresponding gene GTF.

### Statistical analysis

Differential gene expression analysis of scRNAseq data was performed using normalized gene expression counts and the Wilcoxon rank-sum test in the Seurat FindMarkers() function. A log_2_ fold change threshold for gene expression changes of 0.5 and min.pct of 0.25 was used for all comparisons, and a Bonferroni correction was used to control for False Discovery Rate (FDR). A corrected *p* value of < 0.05 considered significant in conjunction with the additional filters above. All other statistical analysis was performed using GraphPad Prism 8 Software (GraphPad Software, La Jolla, CA). A P-valueL<L0.05 was considered significant.

## LIST OF SUPPLEMENTARY MATERIAL

**fig. S1**. Relationship between PCR and scRNAseq viral loads, day 2 post challenge

**fig. S2**. Viral load vs inflammation day 2 post infection

**fig. S3**: Identification and quantification of BALF cells by scRNAseq

**fig. S4:** Identification and quantification of BALF cells by scRNAseq following B.1.3.5.1/beta infection

**fig. S5:** Identification and quantification of SARS-CoV-2 B.1.351/beta variant RNA^+^ cells

## ACKNOWLEDGMENTS

This work was partially supported by a cooperative agreement (W81XWH-18-2-0040) between the Henry M. Jackson Foundation for the Advancement of Military Medicine, Inc., and the U.S. Department of Defense (DOD). The views expressed are those of the authors and should not be construed to represent the positions of the U.S. Army, the Department of Defense, or HJF.

## Author contributions

R.S., M.R., and K.F. designed the study. K.V. and T.L. generated data. A.T.W., K.N., H.F., K.F., M.R. D.L.B., J.R.C., and R.S. analyzed and interpreted the data. A.T.W., K.N., and J.R.C wrote the paper with assistance from all coauthors.

## Data and material availability

All data supporting the findings of this study are available within the manuscript or from the corresponding author upon request. Data tables for expression counts and unprocessed raw data from the scRNAseq analysis are deposited in NCBI’s Gene Expression Omnibus and are accessible through GEO accession GSE190913 (USA-WA1/2020 challenge) and GSE190165 (B.1.351/beta challenge).

## REFERENCES

1. E. Dong, H. Du, L. Gardner, An interactive web-based dashboard to track COVID-19 in real time. Lancet Infect Dis 20, 533–534 (2020).

2. S. Richardson et al., Presenting Characteristics, Comorbidities, and Outcomes Among 5700 Patients Hospitalized With COVID-19 in the New York City Area. JAMA 323, 2052–2059 (2020).

3. S. Gupta et al., Factors Associated With Death in Critically Ill Patients With Coronavirus Disease 2019 in the US. JAMA Intern Med 180, 1436–1447 (2020).

4. R. L. Chua et al., COVID-19 severity correlates with airway epithelium-immune cell interactions identified by single-cell analysis. Nat Biotechnol 38, 970–979 (2020).

5. R. A. Grant et al., Circuits between infected macrophages and T cells in SARS-CoV-2 pneumonia. Nature 590, 635–641 (2021).

6. J. Schulte-Schrepping et al., Severe COVID-19 Is Marked by a Dysregulated Myeloid Cell Compartment. Cell 182, 1419–1440 e1423 (2020).

7. A. J. Wilk et al., A single-cell atlas of the peripheral immune response in patients with severe COVID-19. Nat Med 26, 1070–1076 (2020).

8. W. Wen et al., Immune cell profiling of COVID-19 patients in the recovery stage by single-cell sequencing. Cell Discov 6, 31 (2020).

9. A. Silvin et al., Elevated Calprotectin and Abnormal Myeloid Cell Subsets Discriminate Severe from Mild COVID-19. Cell 182, 1401–1418 e1418 (2020).

10. H. Chen et al., SARS-CoV-2 activates lung epithelial cell proinflammatory signaling and leads to immune dysregulation in COVID-19 patients. EBioMedicine 70, 103500 (2021).

11. C. Yao et al., Cell-Type-Specific Immune Dysregulation in Severely Ill COVID-19 Patients. Cell Rep 34, 108590 (2021).

12. J. S. Lee et al., Immunophenotyping of COVID-19 and influenza highlights the role of type I interferons in development of severe COVID-19. Sci Immunol 5, (2020).

13. L. R. Wong, S. Perlman, Immune dysregulation and immunopathology induced by SARS-CoV-2 and related coronaviruses - are we our own worst enemy? Nat Rev Immunol, (2021).

14. E. Speranza et al., Single-cell RNA sequencing reveals SARS-CoV-2 infection dynamics in lungs of African green monkeys. Sci Transl Med 13, (2021).

15. T. N. Hoang et al., Baricitinib treatment resolves lower-airway macrophage inflammation and neutrophil recruitment in SARS-CoV-2-infected rhesus macaques. Cell 184, 460–475 e421 (2021).

16. P. J. Klasse, D. F. Nixon, J. P. Moore, Immunogenicity of clinically relevant SARS-CoV-2 vaccines in nonhuman primates and humans. Sci Adv 7, (2021).

17. J. Yu et al., DNA vaccine protection against SARS-CoV-2 in rhesus macaques. Science 369, 806–811 (2020).

18. N. B. Mercado et al., Single-shot Ad26 vaccine protects against SARS-CoV-2 in rhesus macaques. Nature 586, 583–588 (2020).

19. K. S. Corbett et al., Evaluation of the mRNA-1273 Vaccine against SARS-CoV-2 in Nonhuman Primates. N Engl J Med 383, 1544–1555 (2020).

20. V. J. Munster et al., Respiratory disease in rhesus macaques inoculated with SARS-CoV-2. Nature 585, 268–272 (2020).

21. H. A. D. King et al., Efficacy and breadth of adjuvanted SARS-CoV-2 receptor-binding domain nanoparticle vaccine in macaques. Proc Natl Acad Sci U S A 118, (2021).

22. M. G. Joyce et al., A SARS-CoV-2 ferritin nanoparticle vaccine elicits protective immune responses in nonhuman primates. Sci Transl Med, eabi5735 (2021).

23. J. S. Tregoning, K. E. Flight, S. L. Higham, Z. Wang, B. F. Pierce, Progress of the COVID-19 vaccine effort: viruses, vaccines and variants versus efficacy, effectiveness and escape. Nat Rev Immunol 21, 626–636 (2021).

24. F. P. Polack et al., Safety and Efficacy of the BNT162b2 mRNA Covid-19 Vaccine. N Engl J Med 383, 2603–2615 (2020).

25. L. R. Baden et al., Efficacy and Safety of the mRNA-1273 SARS-CoV-2 Vaccine. N Engl J Med 384, 403–416 (2021).

26. Y. Goldberg et al., Waning Immunity after the BNT162b2 Vaccine in Israel. N Engl J Med 385, e85 (2021).

27. E. G. Levin et al., Waning Immune Humoral Response to BNT162b2 Covid-19 Vaccine over 6 Months. N Engl J Med 385, e84 (2021).

28. A. Choi et al., Safety and immunogenicity of SARS-CoV-2 variant mRNA vaccine boosters in healthy adults: an interim analysis. Nat Med 27, 2025–2031 (2021).

29. K. S. Corbett et al., Immune correlates of protection by mRNA-1273 vaccine against SARS-CoV-2 in nonhuman primates. Science 373, eabj0299 (2021).

30. K. S. Corbett et al., Evaluation of mRNA-1273 against SARS-CoV-2 B.1.351 Infection in Nonhuman Primates. bioRxiv, (2021).

31. E. Madissoon et al., scRNA-seq assessment of the human lung, spleen, and esophagus tissue stability after cold preservation. Genome Biol 21, 1 (2019).

32. S. Feng et al., Correlates of protection against symptomatic and asymptomatic SARS-CoV-2 infection. Nat Med 27, 2032–2040 (2021).

33. P. Bost et al., Host-Viral Infection Maps Reveal Signatures of Severe COVID-19 Patients. Cell 181, 1475–1488 e1412 (2020).

34. S. Lukassen et al., SARS-CoV-2 receptor ACE2 and TMPRSS2 are primarily expressed in bronchial transient secretory cells. EMBO J 39, e105114 (2020).

35. S. L. Bixler et al., Aerosol Exposure of Cynomolgus Macaques to SARS-CoV-2 Results in More Severe Pathology than Existing Models. bioRxiv, 2021.2004.2027.441510 (2021).

36. J. Pallesen et al., Immunogenicity and structures of a rationally designed prefusion MERS-CoV spike antigen. Proc Natl Acad Sci U S A 114, E7348–E7357 (2017).

37. D. Wrapp et al., Cryo-EM structure of the 2019-nCoV spike in the prefusion conformation. Science 367, 1260–1263 (2020).

38. K. S. Corbett et al., SARS-CoV-2 mRNA vaccine design enabled by prototype pathogen preparedness. Nature 586, 567–571 (2020).

39. K. J. Hassett et al., Optimization of Lipid Nanoparticles for Intramuscular Administration of mRNA Vaccines. Mol Ther Nucleic Acids 15, 1–11 (2019).

40. J. R. Francica et al., Vaccination with SARS-CoV-2 Spike Protein and AS03 Adjuvant Induces Rapid Anamnestic Antibodies in the Lung and Protects Against Virus Challenge in Nonhuman Primates. bioRxiv, (2021).

41. L. A. Jackson et al., An mRNA Vaccine against SARS-CoV-2 - Preliminary Report. N Engl J Med 383, 1920–1931 (2020).

42. A. Vanderheiden et al., Development of a Rapid Focus Reduction Neutralization Test Assay for Measuring SARS-CoV-2 Neutralizing Antibodies. Curr Protoc Immunol 131, e116 (2020).

43. L. C. Katzelnick et al., Viridot: An automated virus plaque (immunofocus) counter for the measurement of serological neutralizing responses with application to dengue virus. PLoS Negl Trop Dis 12, e0006862 (2018).

44. G. X. Zheng et al., Massively parallel digital transcriptional profiling of single cells. Nat Commun 8, 14049 (2017).

45. T. Stuart et al., Comprehensive Integration of Single-Cell Data. Cell 177, 1888–1902 e1821 (2019).

46. A. Butler, P. Hoffman, P. Smibert, E. Papalexi, R. Satija, Integrating single-cell transcriptomic data across different conditions, technologies, and species. Nat Biotechnol 36, 411–420 (2018).

47. C. Hafemeister, R. Satija, Normalization and variance stabilization of single-cell RNA-seq data using regularized negative binomial regression. Genome Biol 20, 296 (2019).

