## Supplemental figures for "mRNA-1273 vaccination protects against SARS-CoV-2 elicited lung inflammation in non-human primates"

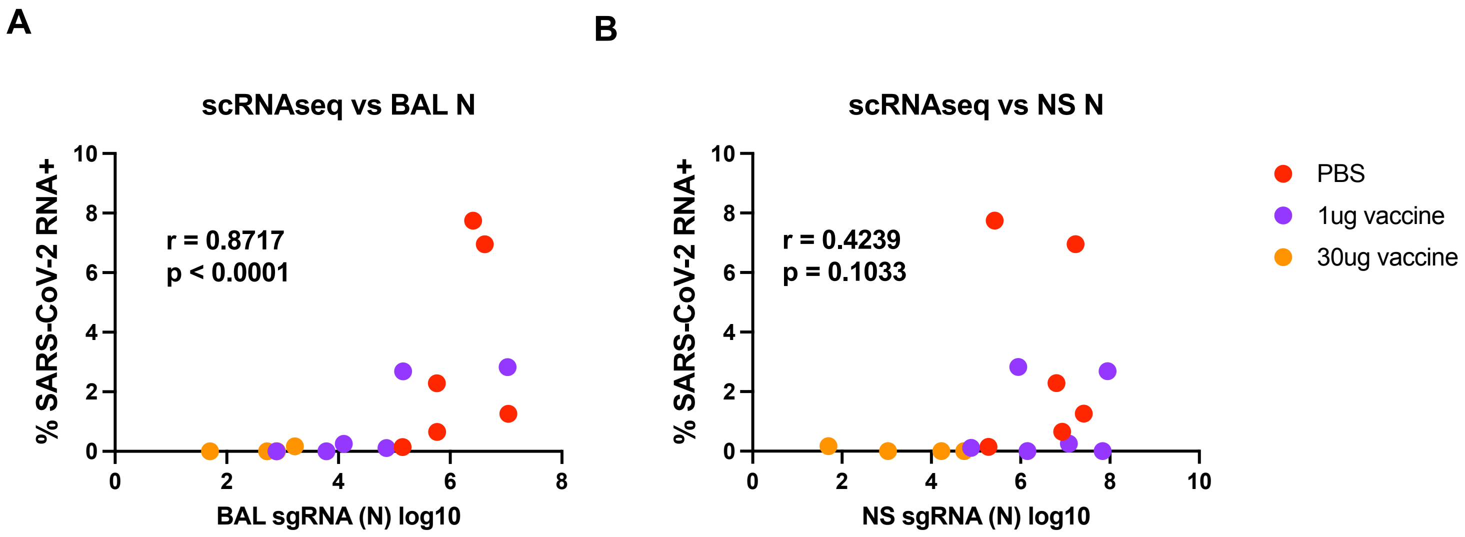


**Fig. S1:** Relationship between PCR and scRNAseq viral loads, day 2 post challenge

A. Relationship between BAL RNA load (N) and frequency of SARS-CoV-2 RNA+ cells
B. Relationship between NS RNA load (N) and frequency of SARS-CoV-2 RNA+ cells

Spearman correlation

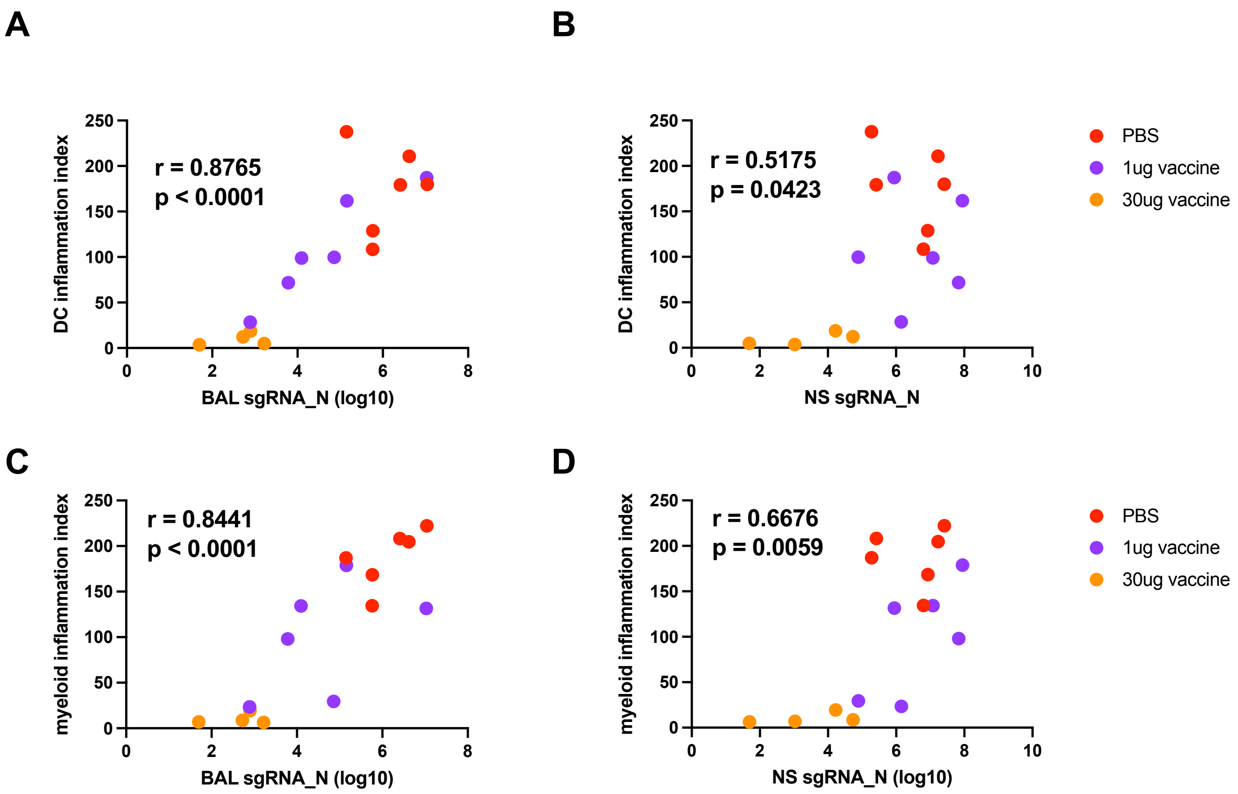


**Fig. S2:** Viral load vs inflammation <u>day 2 post infection</u>

A. DC Inflammation index score vs <u>BAL</u> sgRNA (N gene)
B. DC Inflammation index score vs <u>NS</u> sgRNA (N gene)
C. Macrophage Inflammation index score vs <u>BAL</u> sgRNA (N gene)
D. Macrophage Inflammation index score vs <u>NS</u> sgRNA (N gene)

Spearman correlation

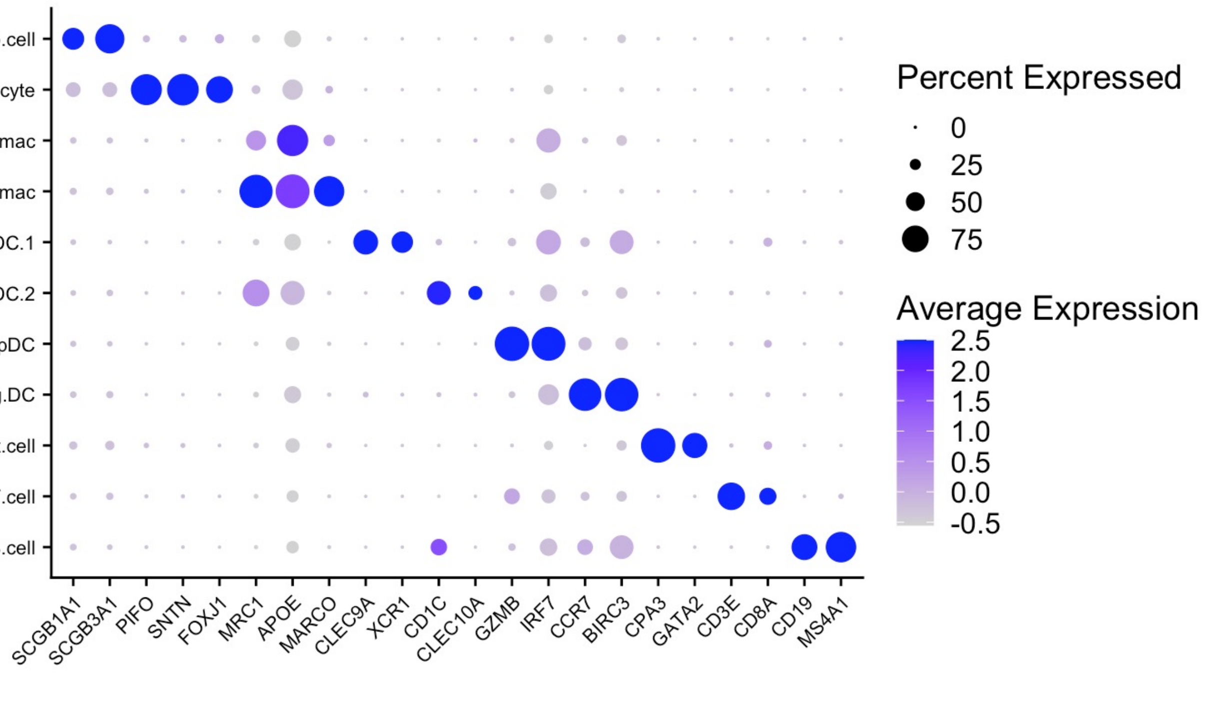


**Fig. S3:** Identification and quantification of BALF cells by scRNAseq

Expression of key linage specific genes in all annotated cell types in BALF collected post SARS-CoV-2 strain B.1.351/beta challenge

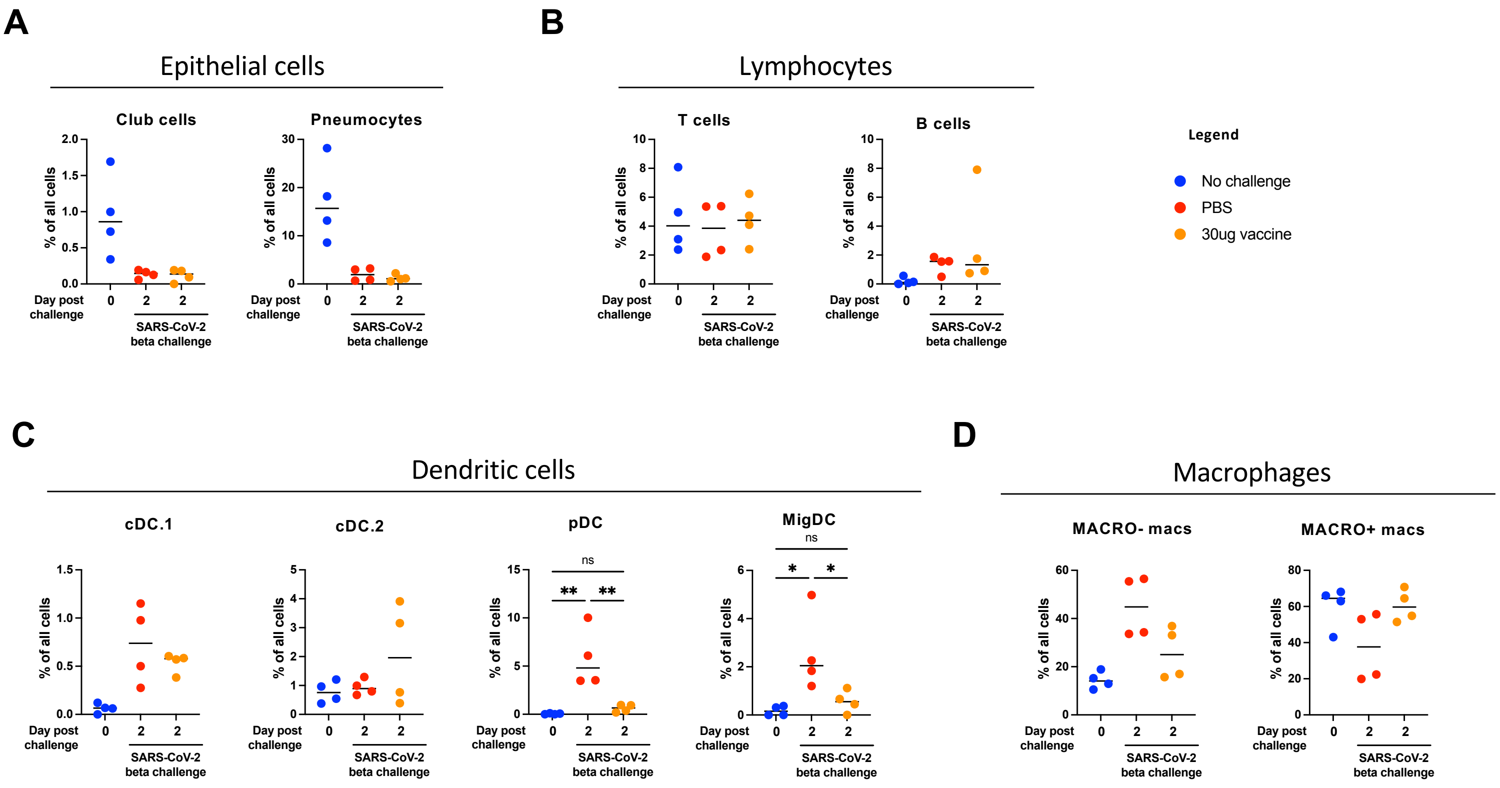


**Fig. S4:** Identification and quantification of BALF cells by scRNAseq following B.1.351/beta infection

A) Frequency of epithelial cell populations
B) Frequency of lymphocyte cell populations
C) Frequency of dendritic cell populations
D) Frequency of macrophage populations

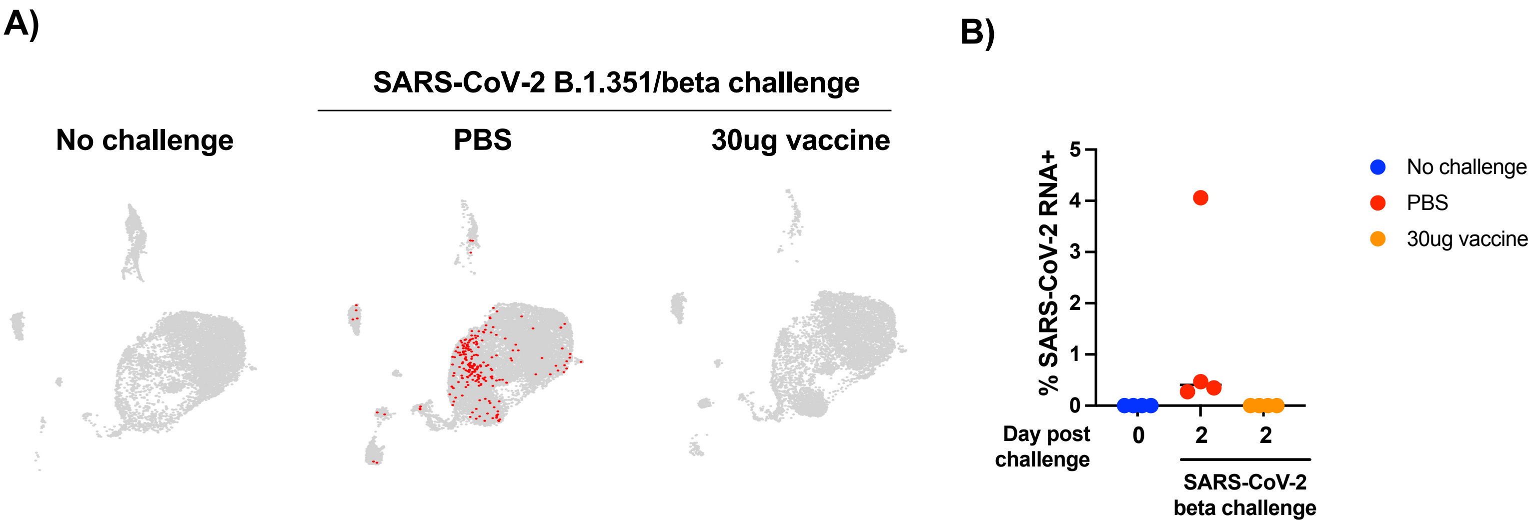


**Fig. S5:** Identification and quantification of SARS-CoV-2 B.1.351/beta variant RNA$^{+}$ cells

A) Location of SARS-CoV-2 RNA$^{+}$ cells

B) Frequency of SARS-CoV-2 RNA$^{+}$ cell
